# Long-lasting effects of transient, perinatal fluoxetine exposure on cell proliferation in the dentate gyrus of mice

**DOI:** 10.1101/778951

**Authors:** Simon C. Spanswick, Michael J. Chrusch, Veronika Kiryanova, Richard H. Dyck

**Affiliations:** Department of Cell Biology and Anatomy, University of Calgary, 3330 Hospital Drive, Calgary, Alberta, T2N 4N1; Hotchkiss Brain Institute, University of Calgary, 3330 Hospital Drive, Calgary, Alberta, T2N 4N1, CANADA; Department of Neuroscience, University of Calgary, 3330 Hospital Drive, Calgary, Alberta, T2N 4N1; Department of Psychology, University of Calgary, 2500 University Drive, Calgary, Alberta, T2N 1N4, CANADA

**Keywords:** Hippocampus, Neurogenesis, Serotonin, SSRI

## Abstract

In the adult mammalian brain, up-regulation of serotonin via the selective serotonin reuptake inhibitor fluoxetine increases hippocampal neurogenesis. However, research assessing the long-term effects of modulating serotonin during the developmental period on hippocampal neurogenesis, is sparse. Here we evaluated hippocampal neurogenesis early (postnatal day 12), and later in life (postnatal day 60), in the offspring of mouse dams that were administered fluoxetine in their drinking water from embryonic day 15 (E15) through postnatal day 12 (P12). Fluoxetine-exposed mice had significantly higher levels of neuronal proliferation at P12, and P60, despite cessation of fluoxetine on P12. These effects were limited to proliferation, as survival of postnatal-born hippocampal neurons was unaltered. Mice exposed to fluoxetine also showed significantly higher levels of cell death, suggesting that homeostatic mechanisms present within the hippocampus may limit integration of adult-born neurons into the existing neuronal network. These findings demonstrate modulation of serotonin during development may be sufficient to induce long-lasting changes in hippocampal neurogenesis.

---

Adult hippocampal neurogenesis is now a well-described phenomenon. The processes of cell proliferation, migration, survival, and integration into an existing brain network have received great attention in the relatively recent past. We now understand that precursor cells within the sub-granular zone of the adult hippocampus give rise to neurons that functionally integrate into the existing hippocampal network [1, 2], eventually becoming indistinguishable from their mature counterparts [3]. A growing body of literature now supports a role for adult-born neurons in behaviour [4, 5, 6, 7].

In addition to the description of adult hippocampal neurogenesis, a vast array of modulatory factors have been discovered, including pharmacological agents [8], aging [9], stress [10], diet [11], exercise [12], and environmental enrichment [13], to name a few. One of the earliest reports of pharmacological modulation of neurogenesis in the adult hippocampus involved chronic administration of the selective serotonin reuptake inhibitor (SSRI) fluoxetine. Malberg and colleagues [14] were the first to report that chronic (14 or 28 day) administration of fluoxetine resulted in an increase of proliferation and survival of adult-born neurons in the DG of adult rats, potentially providing a mechanism by which SSRIs exert their antidepressant effects. Subsequent research demonstrated that fluoxetine specifically targets a class of early progenitor cells within the DG granule cell layer of mice [15], potentially through 5-HT1A/5-HT2A receptor-mediated pathways [16]. Additional support for the role of serotonin in modulating adult neurogenesis comes from lesion studies that note a significant reduction in adult neurogenesis as a result of damaging input fibers from the median raphe [17].Furthermore, knockout mice lacking the 5-HT1A receptor do not show the expected increase in neurogenesis associated with chronic fluoxetine administration [18].The available evidence points to serotonin as an important factor in the regulation of neurogenesis within the adult hippocampus.

Despite a wealth of information regarding the effects of modulating serotonin during adulthood on neurogenesis in the HPC [19], data regarding the same phenomenon during development is limited, with existing research typically assessing the effects of fluoxetine on varying pathologies, most of which report a mitigation of pathology-induced impairments in hippocampal neurogenesis [20, 21, 22]. Given the importance of serotonin during neurodevelopment [23], and the well-documented effects of serotonin up-regulation via fluoxetine administration on the adult hippocampus [14, 18, 24, 25, 26, 27], we sought to investigate the effects of increasing serotonin via fluoxetine administration on neurogenesis in the hippocampus of C57BL/6 mice early in life. In addition, we assessed neurogenesis and cell death during adulthood, long after the cessation of fluoxetine administration.

## Materials and Methods

### Mice and Fluoxetine Administration

All experimental procedures were approved by the University of Calgary Animal Care Committee, and were performed in accordance with guidelines set by the Canadian Council on Animal Care. Mice were housed in a 12-hour light/dark cycle and had *ad libitum* access to food and water throughout the duration of the experiment. FLX mice were generated by pairing C57BL/6 breeders for 4 days, with embryonic day 0 (E0) determined by the presence of a vaginal plug. C57BL/6 dams were administered fluoxetine (Sigma) at 25 mg/kg/day via drinking water from E15 to P12, as has been described previously [28]. Control mice were weighed concurrently with fluoxetine-treated dams to eliminate handling effects, but did not receive fluoxetine in their drinking water. Date of birth for mice pups was considered P0.

Two groups of pups were sacrificed at P12, (a FLX group n = 5, and a corresponding control group, n = 6). All remaining pups were weaned at P21 and housed in groups of two or three until P60, upon which brain tissue was harvested (see Figure 1 for experimental timeline). A FLX (n = 5) and control group (n = 6) received 3 intraperitoneal injections of bromodeoxyuridine (BrdU, 50 mg/kg; Sigma-Aldrich) on P12, spaced at 8-hour intervals; these mice were weaned and housed until perfusion at P60. Thus, tissue was harvested from four groups of mice at P60; two groups that received BrdU at P12, and an additional two groups of mice used to assess proliferation in adulthood (control, n = 9; FLX, n = 8). To mitigate litter effects, groups of mice were randomly drawn from multiple litters, regardless of sex.

**Figure 1.**
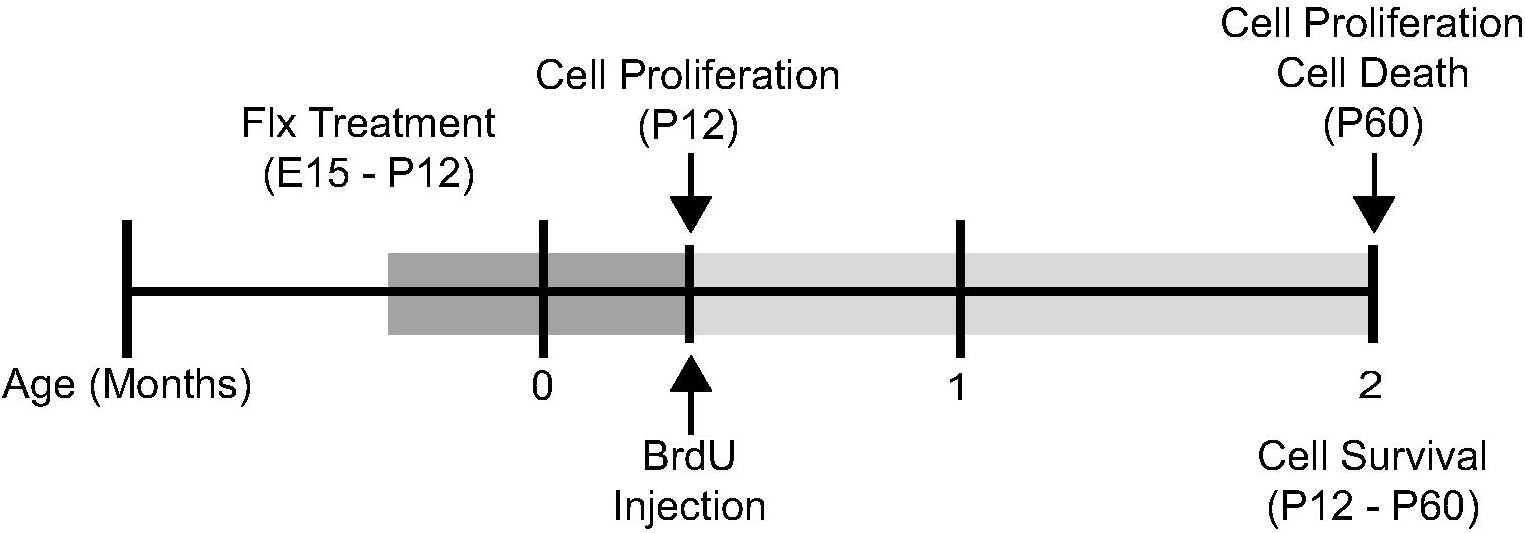
Experimental timeline. C57BL/6 mice were exposed to fluoxetine (Flx) from embryonic day 15 (E15) to postnatal day 12 (P12). Cell proliferation was assessed at P12, and at P60, in separate groups of mice, cell death was also assessed at P60. A single group of mice received injections of BrdU at P12, and were assessed for cell survival at P60.

### Tissue Preparation and Imunnohistochemistry

Mice were injected intraperitoneally with an overdose of sodium pentobarbital (400 mg/kg), and perfused transcardially with 0.1M phosphate buffered saline (PBS), followed by a solution of 4% paraformaldehyde in PBS (PFA). Brains were then extracted, post-fixed in PFA for 24 hours, and then stored in a 30% sucrose/PBS solution. All brains were sectioned in the coronal plane on a freezing sliding microtome (American Optical, model #860; Buffalo, NY, USA) at a thickness of 40 µm, employing a section-sampling fraction of 1/4 for P12 mice, and 1/6 for P60 mice.

To assess proliferation, a single series of tissue from P12 and P60 mice was labeled for the endogenous cell cycle marker Ki67 using methodology that has been described previously [29]. The antibodies used were rabbit anti-Ki67 (1:1000, Novacastra), goat anti-rabbit biotin (1:500, Jackson ImmunoResearch), and streptadivin 594 (1:500, Molecular Probes). The nuclear marker 4’,6’-diamidino-2-phenylindole (DAPI, Sigma, 1:1000) was added as a counterstain in the tertiary step. Sections were then slide-mounted, coverslipped with fluorescent mounting media, and stored at 4°C in the dark until quantification.

To assess survival, a single series of brain tissue sections was taken from P60 mice that had received BrdU injections on P12. Prior to immunofluorescent detection of BrdU, tissue sections were denatured in 2 M HCl, and subsequently washed for 90 minutes with PBS. The method used to visualize BrdU employed the following antibodies: rat anti-BrdU (1:200, Abd Serotec), goat anti-rat biotin (1:500, Jackson ImmunoResearch), and streptavidin 594 (1:500, Molecular Probes). As above, DAPI was added as a counterstain to the tertiary incubation. An additional series was labeled with BrdU/NeuN, in a procedure identical to that discussed above, with the exception of a mouse anti-NeuN antibody (1:2000, Millipore) and the corresponding secondary, goat anti-mouse Cy2, antibody(1:500, Jackson ImmunoResearch).

A final series of tissue from P60 mice was slide-mounted, dehydrated in an ascending series of ethanol steps, and then stained in a 2% Cresyl Violet solution for 15 minutes. The tissue was washed in a destain solution (1% acetic acid in 95% ethanol), and then rinsed in ethanol, before being coverslipped with Permount™ (Fisher Scientific). Slides were stored flat, and allowed to dry prior to analysis.

### Cell Quantification and Volume Estimation

The total number of immune-positive cells was counted in every section within a single series, excluding those within the uppermost focal plane, to minimize edge artifacts [30]. Only those cells that were located in the DG subgranular zone, or granule cell layer were counted. Quantification of cells in P12 mouse brain tissue was performed under a 100X/1.3 objective, whereas P60 mouse brain tissue was quantified under a 63X/1.4 objective, both of which were attached to a Zeiss Axioskop 2 microscope. The number of cells counted in each case was multiplied by the inverse of the appropriate section-sampling fraction to obtain an estimate of total number of BrdU or Ki67 positive cells per DG.

The volume of the DG granule cell layer was calculated employing the Cavalieri method [31]. Images from a DAPI labeled series were assessed using ImageJ software (http://rsb.info.nih.gov/ij/) in a similar fashion to that which has been described before [32]. Co-localization of BrdU and NeuN was determined in at least 50 cells across the rostral-caudal extent of the DG of each mouse. A cell was deemed positive for BrdU/NeuN if a nuclear signal from both the 488 nm and 561 nm lasers co-localized in the x-, y-, and z-axes. The total number of total BrdU/NeuN positive cells was estimated by multiplying the percentage of BrdU positive cells that also were immuno-reactive for NeuN by the estimate of the total number of BrdU positive cells determined earlier.

Utilizing the methodology described above, pyknotic nuclei were counted in the subgranular zone and granule cell layer across the entire rostral caudal extent of the DG. Pyknosis was operationally defined as condensation of chromatin as evidenced by dark, localized staining with Cresyl Violet. The number of cells deemed pyknotic was multiplied by the inverse of the section-sampling fraction (1/6) to obtain a total number estimate.

### Statistical Analysis

All statistical analysis was performed with SPSS^®^ Statistics (Version 19, IBM^®^), and group means were compared using ANOVA, followed by appropriate post-hoc tests when necessary. All group means are expressed as ± SEM, and a p value of ≤ 0.05 was considered as statistically significant.

## Results

### Cell Proliferation

A two-way ANOVA (age by treatment group) revealed a significant main effect of treatment (control versus fluoxetine-exposed mice (FLX), F (1,24) = 23.93, p = 0.000). The control group had a mean of 5545 ± 114 Ki67 positive cells located within the subgranular zone and granule cell layer of the DG, whereas FLX mice had significantly more, with a group mean of 6368 ± 124 (Figure 2A and 2B).

**Figure 2.**
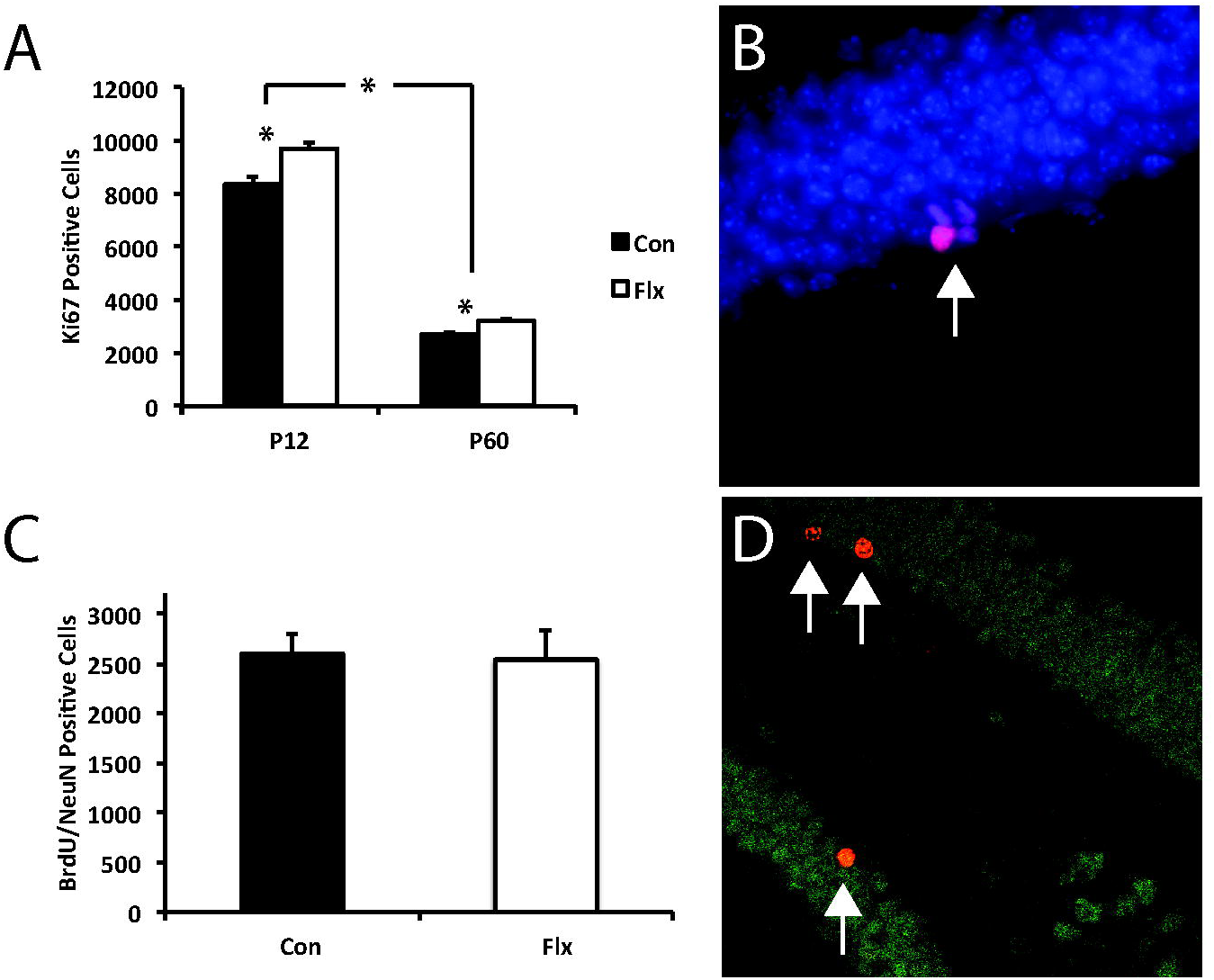
(A) The number of Ki67 positive cells in the subgranular zone and granule cell layer of the dentate gyrus was significantly elevated in fluoxetine (Flx) treated mice at P12 and P60. There was also a significant age-related decline in the number of proliferating cells. * denotes significance, p < 0.05. (B) A cluster of a Ki67 positive cells (red), co-localized with DAPI (blue) within the granule cell layer of the dentate gyrus. (C) Differentiation into a neuronal phenotype was unaltered in Flx treated mice, as measured by BrdU/NeuN colocalization. (D) Cells co-labeled with BrdU (red) and NeuN (green) appear gold, as indicated by arrows.

A significant main effect of age was also apparent (F (1,24) = 1290.07, p = 0.000), with postnatal day 12 (P12) mice having significantly more Ki67-immunoreactive cells than postnatal day 60 (P60) mice (group means 8977 ± 131, and 2936 ± 105, respectively). Unexpectedly, the interaction between age and treatment was also significant (F (1,24) = 4.64, p = 0.042), and as such, post-hoc ANOVAs comparing the treatment conditions at each age were performed utilizing the Bonferroni correction factor (0.05/2 = 0.025).

Post-hoc analysis performed at P12 revealed a significant difference between controls and FLX mice in regards to the number of Ki67 positive cells present, F (1,9) = 11.84, p = 0.007. The control group had a mean of 8385 ± 274 Ki67 positive cells, whereas FLX mice had a group mean of 9570 ± 165 Ki67 cells. The post-hoc analysis performed on the P60 groups was also significant, F (1,15) = 8.45, p = 0.011. Group means were 2706 ± 92 for controls, and 3167 ± 132 for FLX mice. Further post-hoc analysis was performed, assessing each treatment group across the two time points (P12 and P60), applying the same correction factor as mentioned previously. Control animals had significantly fewer Ki67-immunoreactive cells at P60 compared to P12, F (1,13) = 525.75, p < 0.001 (2706 ± 92 vs. 8385 ± 274 respectively). FLX mice also had significantly fewer Ki67 positive cells at P60 than P12, F (1,11) = 848.97, p < 0.001 (9570 ± 165 vs. 3167 ± 132 respectively).

### Cell Survival

Analysis of BrdU-positive cells at P60 revealed no significant difference between controls and FLX mice, F (1,9) = 0.20, p = 0.666. Group means were 3257 ± 305 and 3096 ± 328 BrdU-positive cells, for controls and FLX mice, respectively. The number of BrdU-positive cells also immunoreactive for NeuN did not differ significantly between groups (F (1,9) = 0.05, p = 0.825). The control group had a mean of 2595 ± 190 BrdU/NeuN positive cells, and the FLX group had a mean of 2538 ± 290 BrdU/NeuN positive cells (Figure 2C and 2D). This translated to 76.46 ± 1.83 percent of BrdU positive cells also reactive for NeuN in controls, and 81.95 ± 3.77 percent of BrdU positive cells reactive for NeuN in the FLX group.

### Dentate Gyrus Granule Cell Layer Volume Estimation and Cell Death

To compare DG granule cell layer volumes a two-way ANOVA was performed, with age and treatment as the independent variables. A significant main effect of age was evident, F (1,24) = 8.62, p = 0.007, with P60 mice possessing a larger DG granule cell layer than their P12 counterparts (group means 0.924 ± 0.019 mm^3^, and 0.833 ± 0.024 mm^3^, respectively, Figure 3A). There was no significant main effect of treatment group (p = 0.564), or age by treatment group interaction (p = 0.289).

**Figure 3.**
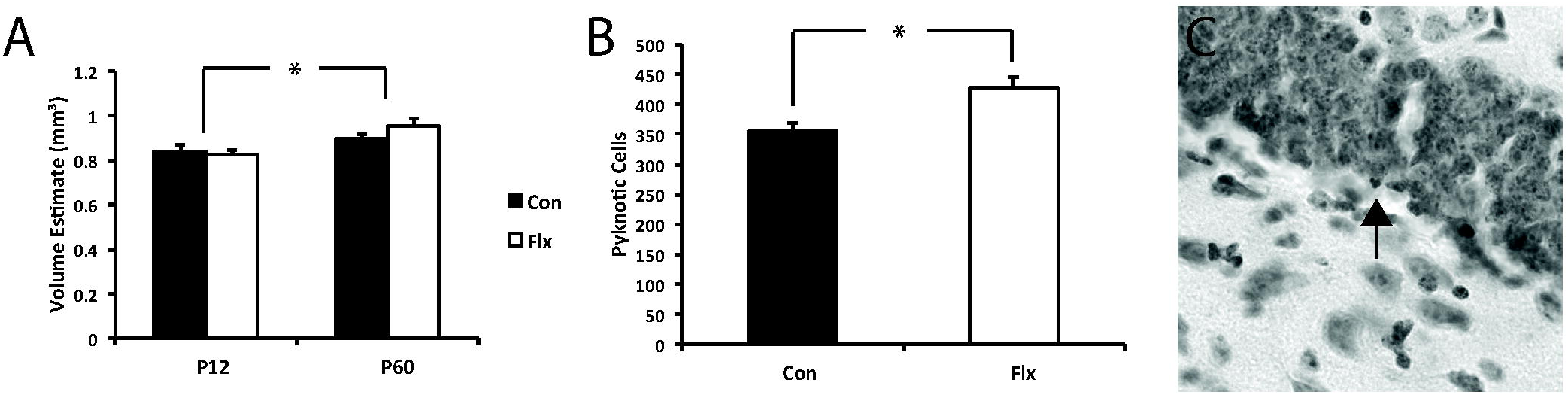
(A) Treatment with Fluoxetine (Flx) had no significant impact on the volume of the dentate gyrus granule cell layer at either P12 or P60. However, there was a significant increase in granule cell layer volume as a function of age. * denotes significance, p < 0.05. (B) Cell death as determined by the presence of pyknotic nuclei was significantly increased in P60 mice exposed to Flx. * denotes significance, p < 0.05. (C) A Cresyl Violet stained section containing a pyknotic nucleus (indicated by arrow) within the dentate gyrus granule cell layer.

A one-way ANOVA revealed a significant difference between control and FLX groups in regards to the number of pyknotic nuclei present in the granule cell layer of the DG, F (1,15) = 10.42, p = 0.006. The control group had a mean of 357 ± 14 pyknotic nuclei, and the FLX group had a mean of 428 ± 17 pyknotic nuclei within the subgranular zone and granule cell layer of the dentate gyrus (Figure 3B and 3C).

## Discussion

Here we show that perinatal up-regulation of serotonergic tone via fluoxetine is sufficient to induce a long-lasting increase in neuronal proliferation in the DG of C57BL/6 mice. Specifically, we observed an increase in the number of Ki67-positive cells in P12 mice that received fluoxetine via the mother from E15 to P12. Importantly, this increase in proliferation was maintained into adulthood (P60) despite cessation of fluoxetine exposure during development (P12). The effects of fluoxetine exposure were limited to proliferation, as mice that were injected with BrdU on P12 did not show enhanced survival of postnatal-born hippocampal granule cells when assessed at P60. This suggests there is an inherent homeostatic mechanism responsible for regulating the basal rate of cellular integration into the existing hippocampal circuitry. Our data showing that there is an increase in cell death within the DG of fluoxetine-exposed mice, and that DG granule cell layer volume was unaltered, lend support to this idea.

The effects of increasing serotoninergic tone via chronic fluoxetine administration on hippocampal neurogenesis within the adult rodent have been well studied. From the initial description of the phenomenon in 2000 by Malberg and colleagues [14], to more recent studies demonstrating the specificity of action at a subset of 5-HT receptors [16], and alterations in vascular dynamics [33], we now possess a clearer understanding of the mechanisms of action by which fluoxetine may up-regulate neurogenesis within the adult hippocampus. Converging evidence suggests that stimulation of the 5-HT2A receptor subtype yields an increase in BDNF mRNA [34], hinting at a mechanism by which long-term increases in hippocampal neurogenesis might be explained. These findings have been supported by more recent research, showing that stimulation of the BDNF Trk receptor is necessary for the observed increase in proliferation associated with fluoxetine exposure [35].

Experiments examining pharmacological up-regulation of serotonin on hippocampal neurogenesis during early development are scant. A small body of evidence suggests that fluoxetine can mitigate the effects of pathology-induced impairments in hippocampal neurogenesis [20, 21, 22]. Here we provide a more direct test of the phenomenon, comparing the effects of increasing serotonin during development in the absence of a disease state.

Curiously, in the absence of pathology, the effects of fluoxetine on hippocampal neurogenesis early in development are varied. Rayen et al. [22] report that in control rats, fluoxetine (administered from P1 – P21) significantly *reduced* cell proliferation (as measured by Ki67) during adolescence (P42). Alternatively, a study by Bianchi et al. [21] found significant increases in both proliferation and survival (1 day and 1 month post-BrdU injection, respectively) in P45 mice that received daily injections of fluoxetine from P3 to P15 (5 to 10 mg/kg, depending upon age). The underlying mechanisms of such effects are not well understood, although the available evidence points towards a BDNF-related pathway [36, 37].

A number of factors may account for the apparently conflicting reports within the literature regarding the effects of perinatal fluoxetine exposure on hippocampal neurogenesis. These include, but are not limited to: dosage of fluoxetine, time-point of assessment, differing immunohistochemical procedures, and method of administration. For example, Bianchi et al. [21] utilized subcutaneous injection in pups, whereas Rayen et al. [22] implanted osmotic minipumps in rat dams on P1. It is conceivable that the stress associated with the aforementioned methods of fluoxetine administration could directly, or indirectly impact ongoing developmental processes, including postnatal neurogenesis. Our method of oral fluoxetine administration via drinking water minimizes stress effects on the dam and her offspring, both of which have been shown to alter hippocampal neurogenesis [38, 39, 40]. In addition, unpublished observations from our laboratory suggest that our method and dose of oral fluoxetine administration is sufficient to produce measurable levels of fluoxetine and its’ metabolite norfluoxetine in the brains of P12 mouse pups [Kiryanova et al., personal communication]. Importantly, these levels fall within the range of those reported in human tissue [41].

In both mice and rats, serotonin transporter protein (5-HTT) is first expressed in the raphe nuclei at E11, with transient expression in non-serotonergic neurons occurring later in development (including limbic regions [42]), even extending beyond birth. We administered fluoxetine to mouse dams from E15 to P12, a time during which transient expression in non-serotonergic neurons of 5-HTT is likely. It may be that pharmacological manipulation of serotonin during this period permanently alters BDNF-related pathways in areas where 5-HTT expression is evident.

Despite significantly increasing proliferation, we found no effects of perinatal fluoxetine on cellular survival 48 days later. These results are consistent with those that report no significant alterations to cell survival after chronic fluoxetine administration in adult animals [43]. Alterations in the latter stages of neurogenesis, such as migration and/or survival often translate to alterations in granule cell layer volume of the DG of rodents in both control [44], and pathological conditions [45, 46]. We report a preferential effect of fluoxetine on proliferation, and no significant change in DG granule cell layer volume in those mice that received fluoxetine, a result consilient with the existing body of literature.

Here we chose to assay the effects of perinatal exposure to fluoxetine in both female and male mice, doing so for two reasons. Evidence shows that there is no sex difference in C57BL/6 mice in terms of hippocampal neurogenesis [47], or volume of the DG granule cell layer [48]. This affords us the ability to combine the sexes, allowing for a significant reduction in the number of litters that were required. Within the clinical setting, the decision to place a pregnant woman on fluoxetine is unlikely to be affected by the sex of her offspring. In addition, our timeline of fluoxetine administration is coincident with the second and third trimester in humans [28], a time during which administration of SSRIs can alter neonatal outcomes [42].

It is possible that fluoxetine exposure altered maternal behaviour and thereby affected the outcomes seen in offspring indirectly. There is evidence to suggest that variability in maternal behaviour can result in long-term changes in central nervous system function in offspring [49, 50]. The data surrounding maternal fluoxetine administration and behaviour is limited, with some studies reporting subtle changes to nursing behaviour [51], while others do not [22, 52].Unpublished data from our laboratory suggests that maternal fluoxetine does not significantly alter the dams’ behaviour toward the pups. As such, the effects of fluoxetine on the mother are likely to be manifest only subtly on the pups’ brain or behaviour. Given the observed increase in anxiety-like behaviors in those studies that report fluoxetine-mediated alterations to maternal behavior [51], they may in fact *attenuate* the increased levels of neurogenesis that we report here [53].

In addition to alterations in proliferation, we provide evidence that cell death is increased in the DG granule cell layer of adult animals exposed to fluoxetine perinatally. We show that the number of pyknotic nuclei is significantly greater in the DG granule cell layer of those adult mice that were exposed to fluoxetine during development than in control animals. Importantly, the increase in cell death is of the same magnitude as the increase in proliferation, strongly implicating a homeostatic mechanism responsible for maintaining normal levels of cellular integration in the DG. It is likely that the increased cell death we observe at P60 is a lifelong homeostatic process in response to increased proliferation. Evidence for regulatory neurogenic mechanisms within the hippocampus have been gleaned from studies assessing neurogenesis in the adult murine brain, both at early [54], and later stages of neurogenesis [55]. Our finding that cell death is increased in those animals that received fluoxetine treatment is congruent with a framework that suggests apoptosis may be the default pathway for regulating the integration of adult-born neurons, unless activity- and experience-dependent factors override them, and promote neuronal survival [56].

In summary, pharmacological up-regulation of serotonin during the perinatal period increases cell proliferation in the DG of adult mice, despite cessation of treatment during the early postnatal period. This increase in proliferation was accompanied by an increase in cell death, providing support for endogenous homeostatic mechanisms that modulate the functional integration of newly born cells into the existing hippocampal neuronal network.

